# Development, characterization and standardization of a nose-only inhalation exposure system for exposure of rabbits to small particle aerosols containing *Francisella tularensis*

**DOI:** 10.1101/570432

**Authors:** Katherine J. O’ Malley, Jennifer D. Bowling, Eileen M. Barry, Karsten R. O. Hazlett, Douglas S. Reed

**Author notes:** address correspondence to Douglas S. Reed, University of Pittsburgh, 9043 Biomedical Science Tower 3, 3051 Fifth Avenue, Pittsburgh, PA 15261. denotes equal contribution.

## Abstract

Inhalation of *Francisella tularensis* (*Ft*) causes pneumonic tularemia in humans, a severe disease with a 30-60% mortality rate. Reproducible delivery of aerosolized virulent bacteria in relevant animal models is essential for evaluating medical countermeasures. Here we developed optimized protocols for infecting New Zealand White (NZW) rabbits with aerosols containing *Ft*. We evaluated relative humidity, aerosol exposure technique, and bacterial culture conditions to optimize spray factor (SF), a central metric of aerosolization. This optimization reduced both inter-and intra-daily variability and were applicable to multiple isolates of *Ft*. Further improvements in the accuracy and precision of the inhaled pathogen dose were achieved through enhanced correlation of bacterial culture OD and CFU. Plethysmograph data collected during exposures found that respiratory function varied considerably between rabbits, was not a function of weight, and did not improve with acclimation to the system. Vaccine Strain (LVS)-vaccinated rabbits were challenged via aerosol with human-virulent *Ft* SCHU S4 that had been cultivated in either Mueller Hinton Broth (MHB) or Brain Heart Infusion (BHI) broth. LVS-vaccinated animals challenged with MHB-SCHU S4 experienced short febrile periods (median: 3.2 days), limited weight loss (< 5%), and longer median survival times (~18 d) that were significantly different than unvaccinated controls. In contrast, LVS-vaccinated rabbits challenged with BHI SCHU S4 experienced longer febrile periods (median: 5.5 days), greater weight loss (> 10%), and median survival times that were not significantly different than unvaccinated controls. These studies highlight the importance of careful characterization and optimization of protocols for aerosol challenge with pathogenic agents.

## INTRODUCTION

*Francisella tularensis* (*Ft*) is a gram-negative coccobacillus that causes a severe zoonotic disease in humans known as tularemia (a.k.a. rabbit fever). Transmission of *Ft* is by direct contact with infected animals or tissues, arthropods, ingestion, or inhalation; tularemia is not typically spread person-to-person. Inhalation of as few as 15 organisms is sufficient to cause disease; *Ft* is readily grown to high concentration in culture and relatively stable in aerosol, properties considered ‘ideal’ in a biological weapon. For these reasons, both the former Soviet Union and United States of America (prior to 1969) included *Ft* in their offensive biological weapons programs (1). There are no licensed vaccines for tularemia; antibiotics are typically given and show good efficacy against most isolates of *Ft*. Considering the high morbidity/mortality and infectivity by aerosol, as well as the possibility of antibiotic resistant strains, *Ft* is a tier 1 select agent, those for which there is the most concern about malicious use (2).

We have previously shown that rabbits are a relevant model of the human disease caused by inhalation of a small particle aerosol containing SCHU S4, the virulent prototype type A strain of *Ft*. Very low doses caused fever, weight loss, dehydration, anorexia, and pneumonia in the rabbits (3). Clinical findings included elevated erythrocyte sedimentation rates, lymphopenia, and thrombocytopenia. Radiographs confirmed the development of bacterial pneumonia and naïve rabbits die within 4-7 days after exposure. The cause of death in the rabbits appeared to be severe septic shock and/or acute respiratory distress syndrome. Additionally, we have shown that attenuated strains of *Ft* can protect rabbits against aerosol challenge with virulent SCHU S4 (4–6). The degree of protection against morbidity and mortality is a function of the attenuated strain used, the number of vaccinations, and the route of vaccination. An aerosol prime/boost vaccination regimen with one particular SCHU S4-derived recombinant strain, S4ΔaroD, protected 66-85% of rabbits against aerosolized SCHU S4 at robust challenge doses (20-200 LD_50_) (4).

Incidence of natural tularemia, particularly for inhalation exposure, is so rare and the bacterium causes such severe disease, that the licensure of vaccines or therapeutics for treating tularemia will only be possible via the FDA’s “Animal Rule” (7, 8). The Animal Rule allows for the use of multiple animal models in ‘pivotal’ efficacy studies when human studies are not ethically or logistically possible. These pivotal studies must be conducted under the auspices of “Good Laboratory Practices” (GLP) which require substantial documentation with an implicit goal of insuring rigor and reproducibility. In more recent guidance, the FDA has created an “Animal Model Qualification Program” which allows animal models to be developed and qualified for use in licensing more than one product (7). Requirements include that the route of challenge must be the same as the probable route of exposure in humans, and that as much as possible, well-characterized pathogens that are minimally-passaged and of human origin should be used. For tularemia, as a bioweapon concern, the route of exposure most likely to be employed in a bioweapon attack is dispersal of the bacterium in a small particle aerosol (≤ 5 µm mass median aerodynamic diameter, MMAD) that can penetrate deep into the alveolar regions of the lung (9).

Most *in vivo* research on tularemia has been with mice, although rats, rabbits, and nonhuman primates (NHP) have also been studied (10). The NHP and rabbit studies have all utilized aerosol delivery for respiratory challenge whereas most of the rodent studies have used intranasal (i.n.) or intratracheal (i.t.) inoculations for respiratory challenge. While i.n. and i.t. methods for inoculation of the respiratory tract are relatively easy to perform and do not require the expensive equipment required for ‘true’ aerosol exposures, substantial differences in deposition have been noted based on methodology and individual operator (11). Penetration into the alveolar region, in particular, is highly variable and it is unclear whether the liquid delivered with the pathogenic agent might alter lung physiology, further complicating matters. Studies using ‘true’ aerosols of *Ft* were employed extensively in the 1950’s, 1960’s, and 1970’s (12–23). There has been renewed interest in *Ft* since the anthrax letters in 2001. In 2012, we showed BHI-grown *Ft* had both higher and more consistent aerosol performance than *Ft* grown in Mueller-Hinton broth (MHB) or Chamberlain’s chemically defined medium (24). Relative humidity (RH) was important, with better performance seen at higher RH. Similar aerosol performance was seen between the Live Vaccine Strain (LVS) and SCHU S4. The improvement in aerosol performance with BHI-grown *Ft* did not appear to impact virulence in naive animals, as assessed by exposure of naïve Balb/c mice to lethal doses of LVS. We report here our efforts to develop standardized protocols for aerosol challenge of rabbits and to demonstrate the reproducibility of these protocols. We have previously shown that rabbits are a relevant outbred model of human tularemia (3), with pulmonary architecture similar to human (25). As part of these efforts, we report here that while the choice of culture media for the aerosolized challenge agent did not affect survival of unvaccinated rabbits, it did affect survival of vaccinated rabbits. These latter observations mirror those reported for intranasal challenge of vaccinated inbred mice (26). The implications of these findings and the impact on conducting pivotral vaccine efficacy studies for tularemia and other high-priority, aerosolized agents will be discussed.

## MATERIALS & METHODS

### Biosafety and Regulatory Information

All work with live *Ft* was conducted at biosafety level-3 (BSL-3) or animal biosafety level-3 (ABSL-3). For respiratory protection, all personnel wore powered air purifying respirators (3M GVP-1 PAPR with L-series bumpcap) or used a class III biological safety cabinet. Vesphene II se (1:128 dilution, Steris Corporation, Erie, PA) was used to disinfect all liquid wastes and surfaces associated with the agent. All solid wastes, used caging, and animal wastes, were steam-sterilized. Animal carcasses were digested via alkaline hydrolysis (Peerless Waste Solutions, Holland, MI). The University of Pittsburgh Regional Biocontainment Laboratory (RBL) is a Registered Entity with the CDC/USDA for work with Tier 1 select agents. Work with recombinant *Ft* strains was approved by the University of Pittsburgh’s Institutional Biosafety Committee.

### Rabbits

Young adult male and female New Zealand White (NZW) rabbits (Robinson Services, Inc.) were housed in the University of Pittsburgh RBL at ABSL-3 for the duration of the studies. All studies were performed under protocols approved by the University of Pittsburgh’s Institutional Animal Care and Use Committee. The University of Pittsburgh is fully accredited by the Association for Assessment and Accreditation of Laboratory Animal Care (AAALAC). Rabbits were monitored at least once daily prior to infection and at least twice daily after infection until clinical signs resolved or the animal became moribund. Prior to vaccination, IPTT-300 temperature/ID chips (BioMedic Data Systems, Seaford, DE) were implanted subcutaneously. Body weight was recorded once in the morning and body temperature was recorded twice daily. Temperature was read using a DAS-7000 reader (BioMedic Data Systems). Rabbits that were determined to be moribund (any of the following clinical signs: weight loss ≥ 20%, body temperature < 34°C, unresponsive to prodding, respiratory distress) were first anesthetized with isoflurane (2-5%) and then euthanized promptly by barbiturate overdose (100 mg/kg sodium pentobarbital given i.v. or i.c.).

### Bacteria

*Ft* LVS or SCHU S4 were originally obtained from Gerald Nau and the Dynport Vaccine Company respectively and were stored as single-passage stocks (3). *Ft* SCHU S4 recombinant derivatives generated in the laboratories of Dr. Eileen Barry and Dr. Wayne Conlan, were shipped to the University of Pittsburgh for these experiments. *Ft* was grown first on Cysteine Heart Agar (CHA) for two days prior to growing overnight in a shaker incubator at 200 RPM, using Brain Heart Infusion (BHI) broth supplemented with ferric pyrophosphate and L-cysteine in baffled, vented polycarbonate Erlenmeyer flasks (3). After the exposures were completed, nebulizer and all-glass impinger (AGI) contents were quantified on CHA.

### Aerosol exposures

Aerosols of *Ft* were conducted inside a class III biological safety cabinet (Baker Co., Sanford, ME) located inside the RBL as previously described using the Biaera AeroMP exposure system (Biaera Technologies, Hagerstown, MD) (3). Supplemental Figure 1 shows a technical schematic of the aerosol system used for these studies. Rabbits were exposed two at a time for 10 minutes in a nose-only exposure chamber (CH Technologies, Westwood, NJ) using a vertical discharge 3-jet Collison nebulizer controlled by the AeroMP (See Supplemental Figure 2 A & B). Aerosol samples were collected in an all-glass impinger containing BHI and antifoam A. These samples were plated on CHA to determine bacterial aerosol concentration. Presented or inhaled dose was determined as described previously, as the product of the aerosol concentration, the minute volume (see below under *Plethysmography*) of the rabbit, and the duration of the aerosol (27).

### Plethysmography

Respiratory function data was collected in real-time during aerosol exposures using Buxco XA or Finepointe software (Data Sciences International, St. Paul, MN) during the exposure (See Supplemental Figure 2C). A pneumotach was mounted on each nose-only tube and calibrated using a 10 cc syringe per the manufacturer’s instructions before the aerosol was started. Minute volume, the mean volume of air inhaled into the lungs per minute, was used to determine inhaled dose. Data was exported from the software into an Excel spreadsheet and averaged for the duration of the exposure.

### Aerosol particle size

Mass median aerodynamic diameter (MMAD) as a measure of particle size was determined using an Aerodynamic Particle Sizer 3321 (TSi, Shoreview, MN) attached to the nose-only exposure chamber; 1 µm fluorescein coated beads in BHI broth were aerosolized as a surrogate for *Ft*. Data was transmitted to the AeroMP and recorded at 2:30 intervals during the exposure.

### Statistical methods

Data was collected and organized using spreadsheets in Microsoft Excel; graphing and statistical analyses were conducted using GraphPad Prism.

## RESULTS

### Environmental data and aerosol particle size

During our initial characterization of the aerosol system used for the rabbit studies, we evaluated the environmental conditions inside the rabbit nose-only (RNO) exposure chamber. Supplemental Table 1 shows the mean, median, standard deviation and coefficient of variation (CV) for relative humidity, temperature, air flow rates and pressure during tularemia aerosols in the RNO. Even without supplemental humidification, a high relative humidity was achieved (75%) with little variability from day to day (CV = 0.12). Even when conditions were drier at the aerosol start, relative humidity increased rapidly once an aerosol was initiated and remained elevated for the remaining aerosol exposures that day (Figure 1A). Aerosol particle size was measured using 1 µm fluorescein beads as a surrogate and the results are shown in Figure 1B. The mass median aerodynamic diameter (MMAD) of the particles (3.05 µm) was larger than what would be expected for a Collision 3-jet nebulizer although the geometric square deviation of 1.77 suggests a relatively homogeneous particle size as is typical for the Collison (28, 29). The larger than expected particle size was likely a function of the short distance between the Collison and the RNO exposure chamber and the high RH. However, 80% of the aerosol particles were less than or equal to 5 µm MMAD so they would still be expected to reach the deep lung of the rabbit respiratory tract. Aerosol performance was compared using ‘spray factor’ (SF), the ratio of the aerosol concentration to the nebulizer concentration. For BHI-grown SCHU S4, SF was higher in the RNO without supplemental humidification compared to BHI-grown SCHU S4 aerosols in a rodent whole-body (RWB) exposure chamber with supplemental humidification (Figure 1C). We also evaluated SCHU S4 grown in MHB and found a similar improvement in SF when using the rabbit nose-only tower without supplemental humidification. The SF achieved with MHB-grown SCHU S4 in the RNO was more than a 1-log improvement over MHB-grown SCHU S4 aerosolized into a RWB chamber with supplemental humidification (4.12×10^−7^ compared to 2.67×10^−8^, respectively) (24). However, the SF for MHB SCHU S4 in the RNO was still significantly lower than that for BHI SCHU S4 in the RNO (*p*<0.0001). A higher SF is indicative of better recovery of the agent from an aerosol and a lower nebulizer concentration required to achieve the same desired dose (9, 30).

**Figure 1.**
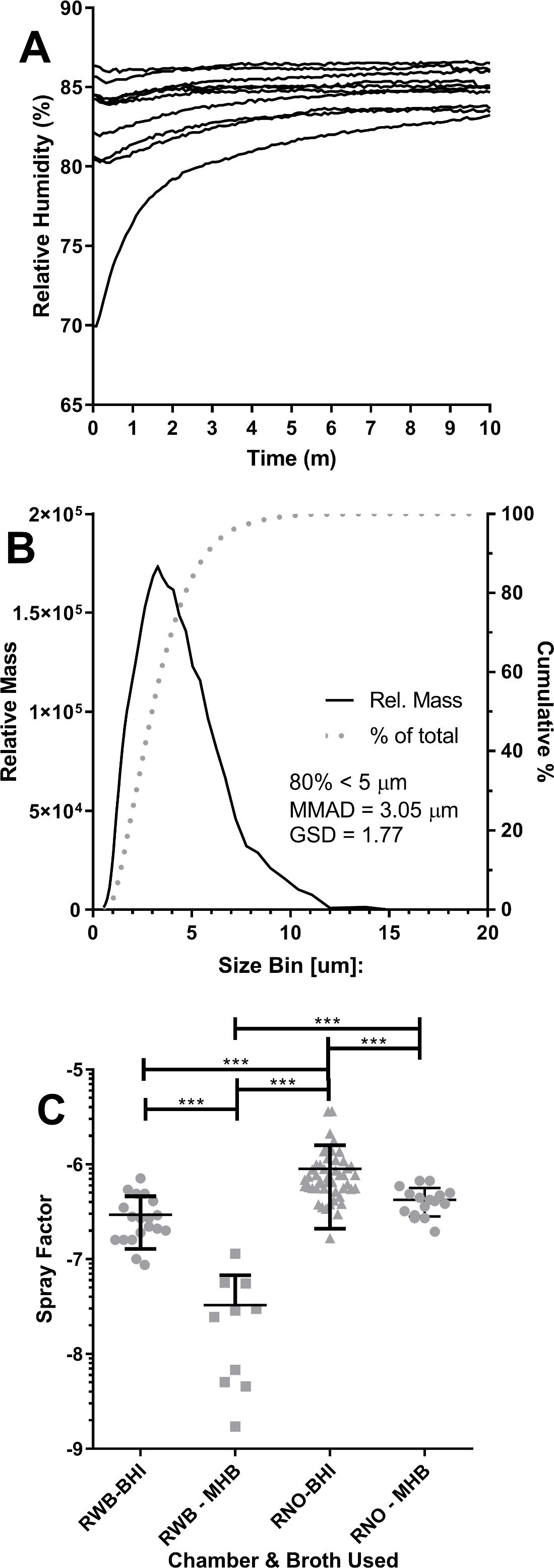
Aerosol performance of the RNO exposure chamber. A) change in relative humidity at 5 second intervals over 10 consecutive 10-minute aerosol exposures in the RNO with *Ft* in BHI. B) Aerosol particle size in the RNO as measured by the APS using 1 µm fluorescein microspheres. The blue line is the relative mass of the particles (left y axis) compared to particle size (x axis); the red line is the cumulative percentage of aerosol particles (right y axis) compared to particle size on the x axis. C) SF for *Ft* SCHU S4 in the RWB and RNO exposure chambers after being grown overnight in either BHI or MHB. Astericks indicate results that were highly significantly different (*p*<0.0001) by t-test with Welch’s correction.

### Reproducibility of system performance

We next sought to evaluate the variability and reproducibility of the RNO for tularemia aerosols. Figure 2A shows the SF for multiple aerosol exposures (‘runs’) conducted on 14 different days using BHI-grown SCHU S4 with the SF mean across all aerosols (9.48×10-7). One-way ANOVA using Brown-Forsythe’s test found that there was a significant difference between means of the different exposure days (*p*=0.0065)(Supplemental Table 2). Tamhane’s T2 multiple comparisons determined that of the 91 comparisons made between days, significant differences were only found in 12 of those comparisons (Supplemental Table 3). We further compared whether SF varied from earlier runs to later runs, evaluating whether samples being kept in an ice bath over the course of several hours affected SF. The results of this comparison are shown in Figure 2B. As the results show, BHI-grown SCHU S4 had a remarkably stable SF across runs over the course of the day, with no significant difference from the mean using a one-way ANOVA with Brown-Forsythe’s test (*p* = 0.7637) as well as Tamhane’s T2 multiple comparisons. We compared whether the SF in the RNO varied between strains and isolates of *Ft* using BHI-grown SCHU S4 and LVS as well as recombinant, attenuated mutants of SCHU S4 (Figure 3). In agreement with what we had previously observed in a RWB chamber, there was no significant difference in SF for any of the isolates by one-way ANOVA with Brown-Forsythe’s test (*p*=0.4076).

**Figure 2.**
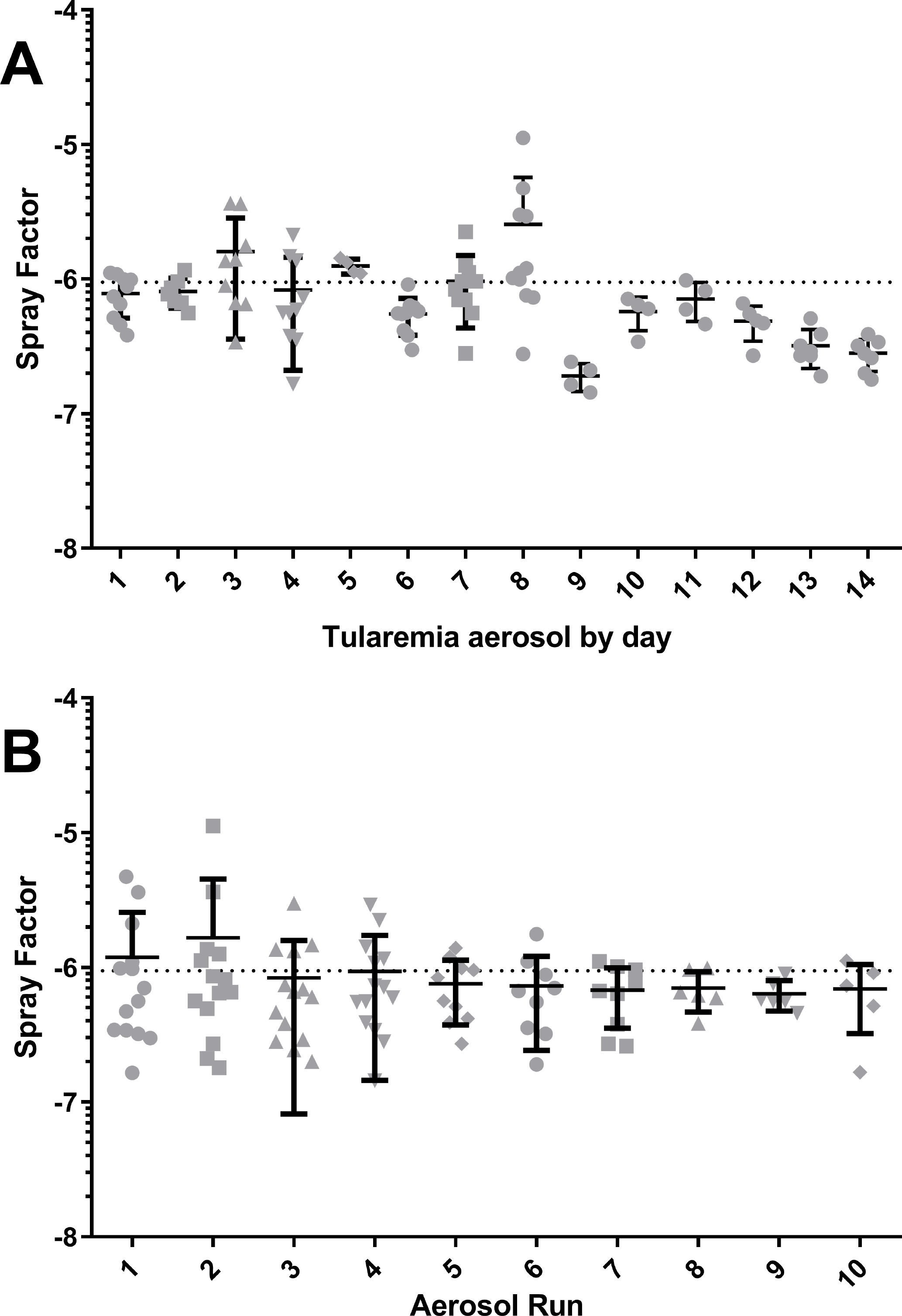
Variation between aerosol exposures of *Ft* SCHU S4. A) Graph shows SF for individual aerosol exposures with the mean and standard deviation for *Ft* aerosols done on 14 different days. Dotted line shows the mean SF (9.48 x 10^−7^) across all SCHU S4 aerosols. B) Graph shows SF for aerosol ‘runs’ over multiple days; values plotted are individual SF for that run from different days with the mean and standard deviation. Dotted line shows the mean SF (9.48 x 10^−7^) across all SCHU S4 aerosols.

**Figure 3.**
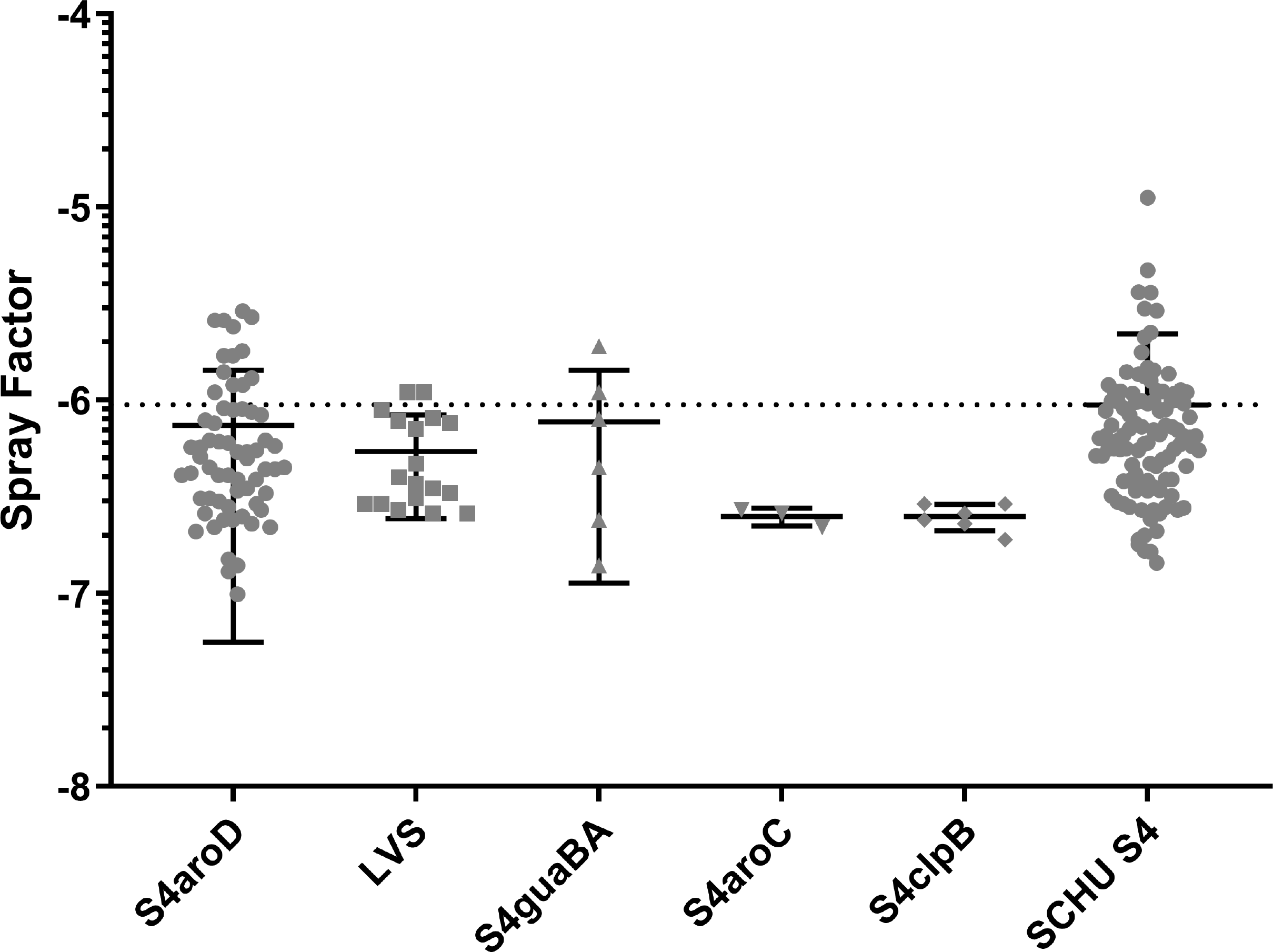
No significant variation in SF between different isolates of *Ft*. SCHU S4, LVS, or recombinant derivative isolates of SCHU S4 were grown in BHI and aerosolized in the RNO. Results shown are the individual SF for different exposures with the mean and standard deviation for each isolate. Dotted line shows the mean SF (9.48 x 10^−7^) for SCHU S4 aerosols.

### Improving accuracy and precision in challenge dose

The most predominant concerns with aerosol exposure of animals to infectious agents is a) achieving the desired target inhaled dose and b) insuring all animals receive comparable doses. Inhaled (a.k.a. the presented dose) is calculated as the product of the aerosol concentration of the agent, the duration of the exposure, and the minute volume (mean volume of air inhaled per minute) of the animal (30). Depending on the agent, environmental parameters and choice of culture media for propagating the agent can affect aerosol concentration, which impacts the inhaled dose. These parameters can be controlled and their effects calculated. In our initial tularemia vaccine study done in rabbits, the mean inhaled challenge dose (11,070 cfu) delivered to the rabbits was 11-fold higher than the desired target dose of 1,000 cfu (Figure 4A). Analysis of the aerosol parameters revealed a 2-fold better than predicted SF (7.8×10^−7^ actual vs 4.4×10^−7^ predicted), but also a nebulizer concentration that was 4-fold higher than desired (9.8×10^5^ actual vs 2.4×10^5^ desired).

**Figure 4.**
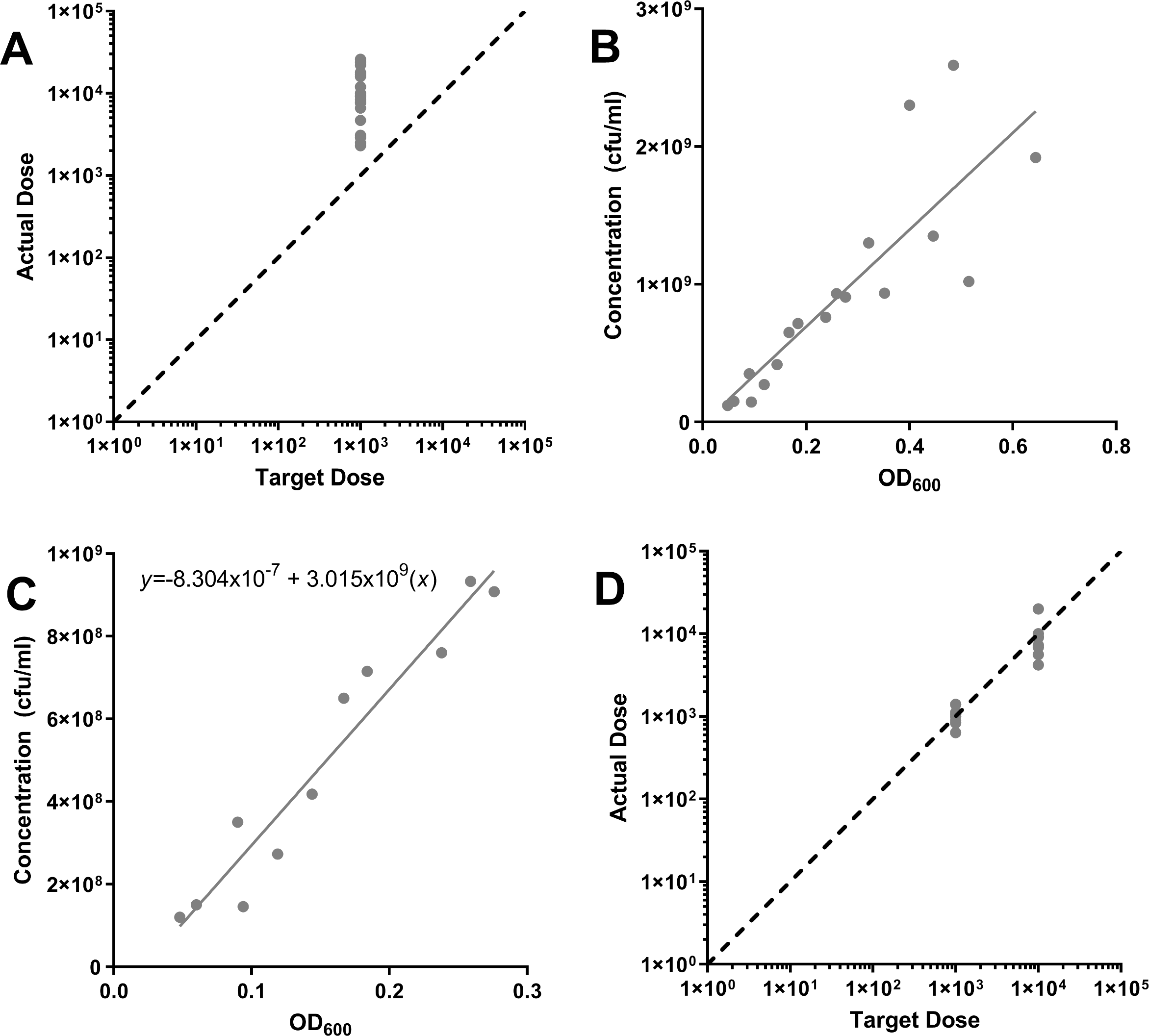
Improvements in inhaled dose based on improved determination of bacterial concentration using OD_600_ within the linear range. A) Aerosol doses in rabbits for the first vaccine study (gray circles are individual rabbits); actual dose is on the y axis while the target dose is on the x axis. B) OD (x axis) and CFU (y axis) readings of different cultures of SCHU S4 grown in BHI used to determine nebulizer concentration for the results shown in A). C) narrowed OD to CFU range to fit linear regression analysis; the resulting formula is shown on the graph. The *r^2^* value was 0.99. D) Aerosol doses in the subsequent vaccine study (gray circles are individual rabbits); actual dose is on the y axis while the target dose is on the x axis.

Although there was variability in the SF from what we had predicted, the data suggested that a more precise estimation of the nebulizer concentration would improve our accuracy in achieving the target dose. Bacterial concentration in broth culture is often estimated using the optical density (OD) of the culture (for *Ft*, OD is read at 600 nm) and comparing that with colony growth on agar. Typically, after an 18-hour culture of SCHU S4 in BHI, the OD_600_ of the culture is ~0.9. In our determination of the OD to CFU ratio, this was beyond that portion of the curve that was linear, which was between 0.05-0.3 (Figure 4B & 4C). We felt that making dilutions of the broth culture to get the OD within that range would produce the desired concentration.

In a subsequent repetition of the vaccine study, rabbits were split into 2 doses; one group was to receive 1,000 cfu while the second was to receive 10,000 cfu. Two-fold dilutions were made from the overnight broth culture until the OD_600_ fell within the linear range, which was used to then back-calculate the broth culture concentration. For the 1,000 cfu target dose group, the actual mean challenge dose was 1,025 cfu while for the 10,000 cfu target dose group the actual mean challenge dose was 9,038 cfu (Figure 4D). For all subsequent vaccine studies, using this refined method for determining the broth culture concentration, challenge doses have averaged 2.6-fold higher than the target dose (Supplemental Figure 2 and Supplemental Table 4).

While this refined methodology for determining bacterial concentration in the overnight broth culture, thereby improving the average actual inhaled challenge dose, it did not improve precision in dosing between animals, as evidenced by the range of inhaled challenge doses (coefficient of variation, Supplemental Table 4). Since the nebulizer concentration and SF were within the range of predicted values and aerosol duration was fixed at 10 minutes, the only remaining variable was the respiratory function of the rabbits. Figure 5 shows the plethysmography data collected in real-time during rabbit aerosol challenges. Predictive formulas such as Guyton’s, Alexander’s, and Bide’s are a function body weight (31–33). As Figure 5A shows, while there is a trend of increased minute volume with weight, for individual rabbits body weight would not be particularly helpful in predicting minute volume. Minute volumes between two rabbits of the same age and weight can vary by more than 1 lpm. When rabbit minute volume was plotted by aerosol exposure days, there was day-to-day variation as shown in Figure 5B and these differences were significant by one-way ANOVA using Brown-Forsythe’s test (*p*<0.0001). In the midst of these studies, we had also evaluated inhalation of small particle aerosols containing attenuated strains as a potential means of delivery for live vaccines (4). Using the data acquired during those studies, we evaluated whether repeated aerosol exposures of rabbits in the nose-only tubes would alter respiratory function (Supplemental Figure 3). There were significant variations by one-way ANOVA in breathing rate (*p*=0.0437), tidal volume (*p*=0.0217), and minute volume (*p*=0.0086) across the three aerosol exposures. However, there was no trend across the three exposures, either positive or negative, therefore acclimation did not appear to alter respiratory function in this circumstance.

**Figure 5.**
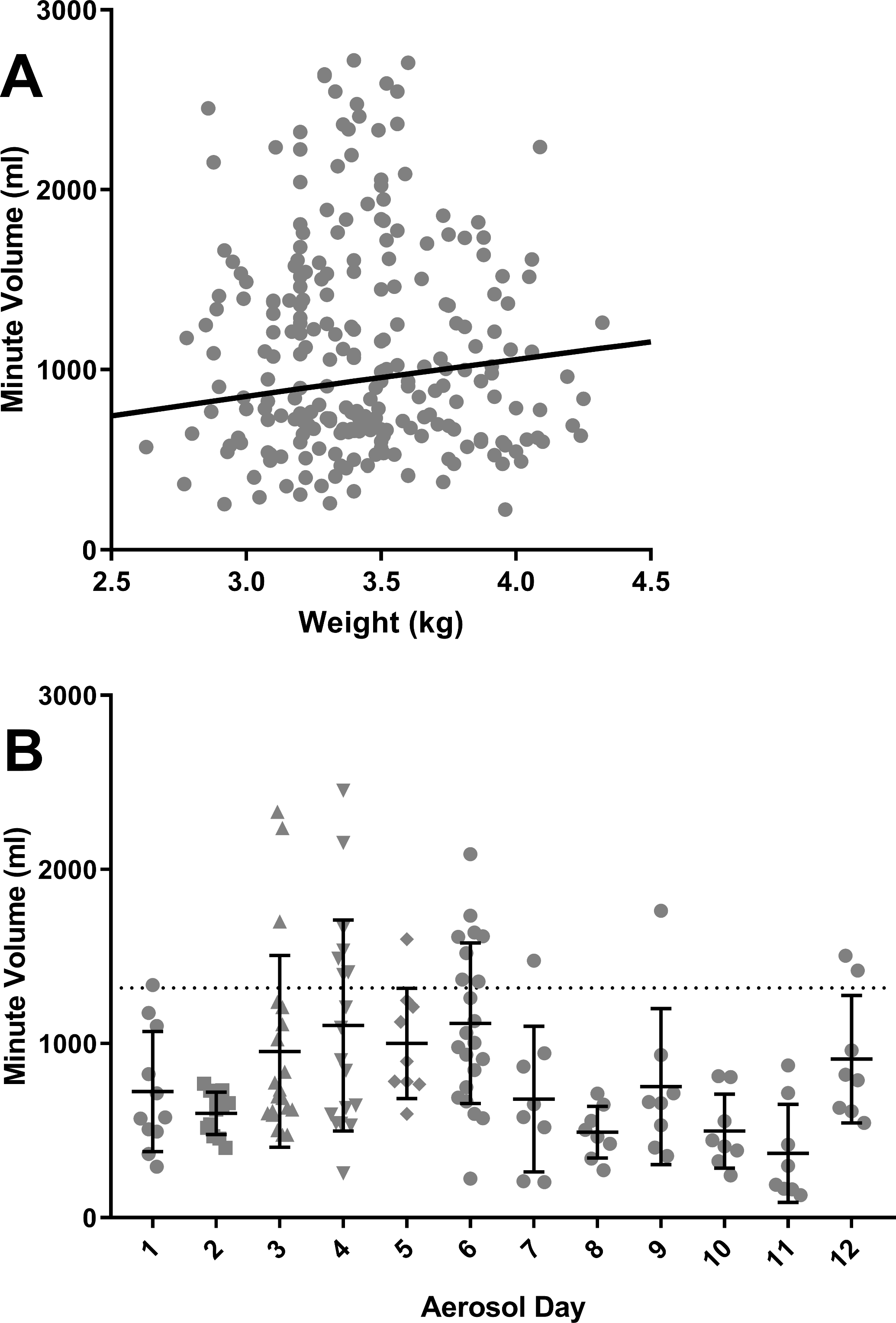
Averaged real-time minute volume in rabbits during aerosol exposures to *Ft*. A) averaged minute volume (y axis) over 10 minute aerosol exposures for individual rabbits (gray circles) compared to baseline weight (x axis); black line is minute volume calculated from body weight using Guyton’s formula. B) averaged minute volume for individual rabbits on the day of exposure. Dotted line is the mean minute volume across all rabbits.

### Challenge broth media impacts survival of LVS-vaccinated rabbits

We recently reported that LVS-vaccinated, inbred mice survived better if challenged with MHB-grown SCHU S4 than BHI-grown SCHU S4 (26). Using the parameters defined above, we sought to evaluate whether the same pattern of vaccine-mediated protection would be seen in aerosol-challenged outbred rabbits. Rabbits were vaccinated with LVS using a prime-boost aerosol vaccination approach, with the boost given 14 days after the prime. Thirty days after the booster vaccination, rabbits were challenged with 500 cfu of SCHU S4 grown in MHB or BHI. LVS-vaccinated rabbits in both groups developed fevers after challenge but those challenged with BHI-grown SCHU S4 had fever responses persist far longer than those challenged with MHB-grown SCHU S4 (Figure 6A). The persistence and severity of the fever response in the BHI-grown SCHU S4 challenged LVS-vaccinated rabbits is further demonstrated in the heat map shown in Figure 6B. Mock-vaccinated rabbits develop fever about the same time regardless of how SCHU S4 is cultured and succumb at roughly the same time (between days 5-6). LVS-vaccinated rabbits in both groups lost weight after challenge but beginning on day 7, weight started to recover in the MHB-challenged rabbits while BHI-challenged rabbits continued to lose weight through day 15 (Figure 6C). At day 20, the last date weight was recorded, the BHI-challenged rabbits had stopped losing weight but had not begun to regain the weight they had lost. Three of six LVS-vaccinated rabbits survived challenge with MHB-grown SCHU S4 while only one of six LVS-vaccinated rabbits survived challenge with BHI-grown SCHU S4 (Figure 6D). The difference in survival for LVS-vaccinated, MHB-SCHU S4 challenged rabbits was statistically significant when compared to historical controls (*p* = 0.0473) by t-test while it was not significant for survival of LVS-vaccinated, BHI-SCHU S4 challenged rabbits when likewise compared to historical controls (*p*=0.0981).

**Figure 6.**
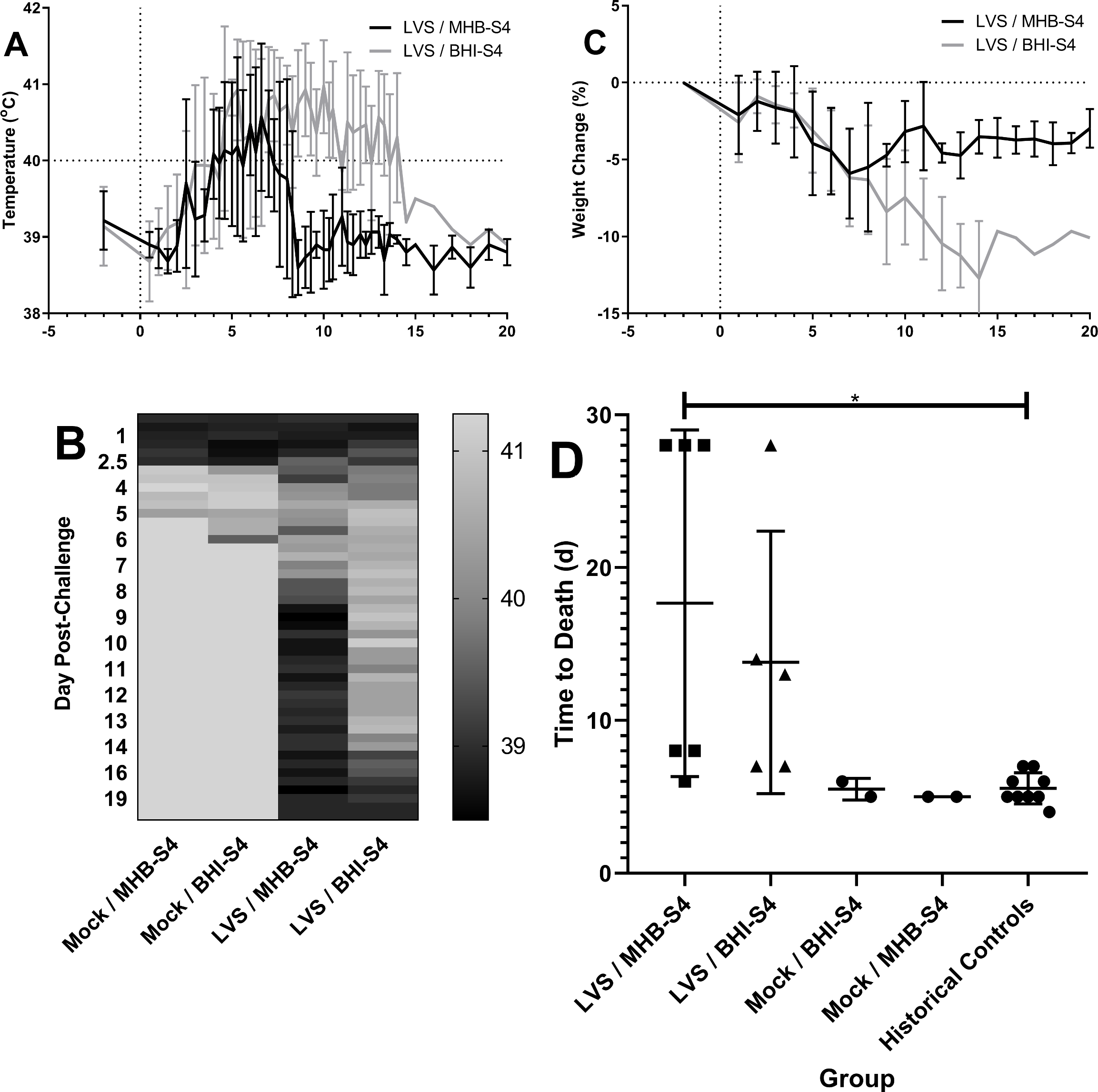
BHI-grown *Ft* is a more rigorous challenge than MHB-grown *Ft* for vaccinated rabbits. Rabbits were vaccinated by aerosol prime-boost delivery of LVS two weeks apart, then challenged 30 days after the boost with ~500 cfu of SCHU S4 grown in MHB (black lines, A & C) or BHI (gray lines, A & C). A) mean body temperature by group with error bars for standard deviation for the first 20 days post-challenge; B) mean body temperature by group as a heat map for the first 20 days post-challenge, including mock-vaccinated groups; C) mean percent weight change by group with error bars for standard deviation for the first 20 days post-challenge; D) time to death in each vaccine and control group, compared to historical mock-vaccinated controls challenged with equivalent doses of BHI-grown SCHU S4. Time-to-death for the LVS-vaccinated, MHB-SCHU S4 challenged rabbits was significantly different from historical controls by one-way ANOVA, *p*<0.05. Time-to-death for the other groups were not significantly different from the historical controls.

## DISCUSSION

We report here our efforts to develop and characterize a standardized, reproducible method for aerosol challenge of rabbits with virulent *Ft*. Using our protocol of growing *Ft* for two days on CHA followed by an overnight broth culture in BHI we found while there is some variability in SF between aerosol exposures there is none between runs on a given exposure day. This consistency will be quite useful in pivotal efficacy studies where standardization and reproducibility will be critical. Refining our ability to pinpoint the concentration of bacteria in our broth culture also allowed us to improve our accuracy in achieving the desired target dose, with only a 2.6-fold difference between actual and target challenge doses over 13 subsequent vaccine studies. The data reported here show the value in developing a well-characterized aerosol challenge system, particularly for vegetative bacteria.

Our protocol did not eliminate all variability, as there was some day-to-day variation in SF as well as other factors. There is the inherent error in current methods used for quantifying bacteria via either spectrophotometry or growth on agar which can add some variability. Our refinement in determining broth culture concentration and subsequent nebulizer concentration improved the accuracy of the mean actual doses obtained in relation to the desired target dose but did not improve precision between animals. Differences in respiratory function between individual animals would also contribute to variability when aerosols are for a fixed time duration (in our case, 10 minutes). Altering the duration of the exposure based on differences in respiratory function would correct for this variability, insuring that all animals breathe in the same volume of experimental air and improving precisions. There are currently two possibilities for adjusting the duration of an aerosol. First, respiratory function can be measured immediately prior to the aerosol and this value put into the aerosol system, adjusting the length of the aerosol. However, this cannot correct for differences in respiratory function that might occur during the aerosol. The second option is to measure respiratory function in real-time and pass that data to the system controlling the aerosol, adjusting the duration of the aerosol on the fly. This has already been demonstrated for NHP using the same Biaera AeroMP system that we use (34), and we are currently evaluating a new version of the Finepoint software (Data Sciences International, St. Paul, MN) that would do the same for rabbits. The only caveat to this is that with the current system, rabbits would have to be done one at a time if aerosol duration were tied to respiratory function. This would double the amount of time required to complete a vaccine challenge compared to our current challenge system.

One argument against using the rabbit for tularemia vaccine studies had been that vaccination with LVS extended time to death but not survival (5, 6, 10, 21). Our own prior studies are included in those findings, including the observation that oral inoculation or inhalation of LVS did extend rabbit time to death more than subcutaneous inoculation (5). This data does fit with what has been reported in humans and monkeys where oral and aerosol LVS vaccination offered better protection against subsequent respiratory challenge (17, 20, 35, 36). The data presented here demonstrate that the apparent ‘failure’ of LVS vaccination in the rabbit model was at least partly due to our use of SCHU S4 grown overnight in BHI broth for aerosolization. MHB is the predominant broth media used for propagation of *Ft*, although MCPH (modified casein partial hydrolysate), TSB (trypticase soy broth), and BHI are also used. To our knowledge, TSB and MCPH for propagation of SCHU S4 have not been as carefully evaluated as have been BHI or MHB. We had originally chosen BHI as *Ft* grown in BHI had a higher, more consistent SF than MHB and growth of *Ft* in BHI did not appear to alter virulence or pathogenicity in naïve animals (24). BHI-grown *Ft* has been shown to more closely resemble *Ft* isolates from infected macrophages than MHB-grown *Ft* (37). We recently reported that LVS-vaccinated mice survived low-dose challenge with MHB-grown SCHU S4 better than after challenge with comparable doses of BHI-grown SCHU S4 (26). Here, we did not see 100% survival of LVS-vaccinated rabbits when challenged with ~500 cfu (~25 LD_50_) MHB-grown SCHU S4, and even those that did survive developed a fever and lost weight post-challenge. However, the severity and duration of both the fever and weight loss seen in LVS-vaccinated rabbits challenged with MHB-grown SCHU S4 was less than that in LVS-vaccinated rabbits challenged with BHI-grown SCHU S4. These data suggest that BHI-grown SCHU S4 is a more rigorous challenge for demonstrating protection in vaccine studies than MHB-grown SCHU S4. Considering that we have previously shown that attenuated derivatives of SCHU S4 outperform LVS and protect rabbits well against both morbidity and mortality at even higher challenge doses of SCHU S4 grown in BHI, this further illustrates the superior performance of these attenuated derivatives as potential vaccine candidates (4–6). The survival of LVS-vaccinated rabbits and consistent/higher SF from the aerosol characterization studies both argue that BHI should be used in pivotal *Ft* efficacy studies to achieve a rigorous, reproducible aerosol challenge sufficient to demonstrate the efficacy of potential vaccine candidates in animals.

## ACKNOWLEDGEMENTS

The research described herein was sponsored by the National Institute of Allergy & Infectious Diseases at the National Institutes of Health, grants U01 AI077909-01, R01 A102966-01A1, AI 123129, and AI 100138

## SUPPLEMENTAL FIGURE LEGENDS

**Supplemental Figure 1. Aerosol system design.** The schematic shows the overall design and flow of the aerosol system used to expose rabbits to *Ft*, including the use of plethysmography to monitor respiratory function during aerosols. The AeroMP is the aerosol management platform from Biaera Technologies (Hagerstown, MD).

**Supplemental Figure 2. Aerosol exposures of rabbits to *Ft*.** Pictures show rabbits in nose-only exposure tubes attached to the tower (top picture), the computer controlling the AeroMP system (middle picture), and an image of plethysmography data being collected during an exposure (bottom picture).

**Supplemental Figure 3. Improved precision and accuracy in challenge doses delivered to rabbits.** Graphs show actual (y axis) vs target (x axis) doses for individual rabbits (gray circles) in four separate vaccine studies after adopting our refinement in determining the concentration of our SCHU S4 broth culture.

**Supplemental Figure 4. No evidence for acclimation of rabbits to aerosol exposures.** Graphs show averaged respiratory function for individual rabbits (gray circles) exposed 1, 2 or 3 times to *Ft*. A) breathing rate, B) tidal volume, C) minute volume in rabbits. There was no significant change in any of the respiratory parameters examined over repeated aerosol exposures.

## SUPPLEMENTAL TABLES

**Supplemental Table 1.**
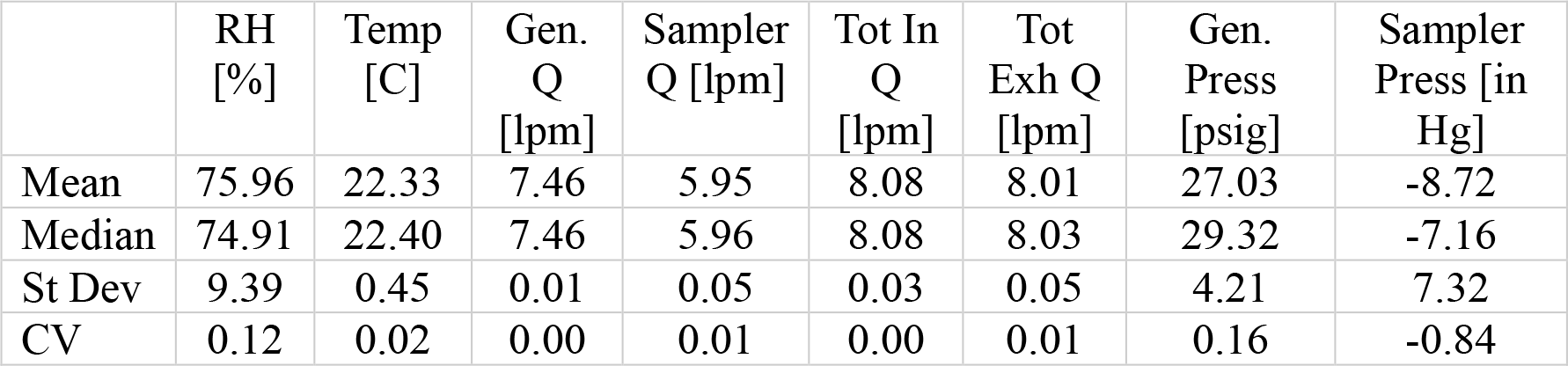
Environmental parameters inside the exposure chamber for aerosol exposures of rabbits to *Ft* grown in BHI. RH = relative percent humidity, Temp = temperature (in Celsius), Gen. Q = generator flow rate, in liters per minute (lpm), Sample Q = flow rate through the all-glass impinger, Tot In Q = total system input flow rate, Tot Exh Q = total exhaust flow rate, Gen Press = generator pressure, in pounds per square inch gauge (psig), Sampler Press = sampler pressure, in millimeters of mercury (Hg).

**Supplemental Table 2.** ANOVA analysis of day-to-day variability in tularemia spray factor.

| Test | Result |
| --- | --- |
| Brown-Forsythe ANOVA test |  |
| F* (DFn, DFd) | 3.936 (13.00, 15.19) |
| P value | 0.0065 |
| P value summary | ** |
| Significant diff. among means (P < 0.05)? | Yes |
| Welch's ANOVA test |  |
| W (DFn, DFd) | 18.89 (13.00, 27.85) |
| P value | <0.0001 |
| P value summary | **** |
| Significant diff. among means (P < 0.05)? | Yes |
| Data summary |  |
| Number of treatments (columns) | 14 |
| Number of values (total) | 102 |

**Supplemental Table 3.** Tamhane’s T2 multiple comparisons analysis of day-to-day variability in tularemia spray factor. Results shown are those that were significantly different, *p*<0.05. Groups = SF days being compared; Mean Diff. = difference between means; Adj. p value = adjusted p value, based on Tamhane’s analysis.

| Groups | Mean Diff. | Adj. p value |
| --- | --- | --- |
| 1 vs. 9 | 5.90E-07 | 0.0044 |
| 1 vs. 13 | 4.60E-07 | 0.0284 |
| 1 vs. 14 | 4.99E-07 | 0.0153 |
| 2 vs. 9 | 6.18E-07 | 0.0105 |
| 2 vs. 13 | 4.89E-07 | 0.0281 |
| 2 vs. 14 | 5.28E-07 | 0.0221 |
| 5 vs. 6 | 6.97E-07 | 0.033 |
| 5 vs. 9 | 1.06E-06 | 0.0427 |
| 5 vs. 12 | 7.59E-07 | 0.0234 |
| 5 vs. 13 | 9.28E-07 | 0.0227 |
| 5 vs. 14 | 9.66E-07 | 0.0397 |
| 6 vs. 9 | 3.60E-07 | 0.0048 |

**Supplemental Table 4.** Difference between target and actual challenge doses for vaccine studies. OD Optimized, N= no, Y=yes; n = number of rabbits challenged; Target = target inhaled dose, in cfu; Actual = averaged actual inhaled dose, in cfu; Fold Difference = difference between actual and target dose; Min = minimum inhaled dose; Max = maximum inhaled dose; CV = coefficient of variation.

| OD Optimized | n | Target | Actual | Fold Difference | Min | Max | CV |
| --- | --- | --- | --- | --- | --- | --- | --- |
| N | 20 | 1,000 | 11,070 | 11.1 | 2,300 | 26,000 | 0.65 |
| Y | 12 | 1,000 | 1,025 | 1.0 | 230 | 4,200 | 1.11 |
| Y | 8 | 10,000 | 9,038 | 0.9 | 2,500 | 55,000 | 2.82 |
| Y | 12 | 1,000 | 1,373 | 1.4 | 636 | 1,397 | 0.16 |
| Y | 6 | 10,000 | 21,800 | 2.2 | 4,200 | 20,000 | 0.22 |
| Y | 8 | 1,000 | 6,051 | 6.1 | 4,203 | 8,777 | 0.24 |
| Y | 20 | 1,000 | 660 | 0.7 | 105 | 2,098 | 0.64 |
| Y | 20 | 1,000 | 713 | 0.7 | 140 | 1,401 | 0.52 |
| Y | 20 | 200 | 333 | 1.7 | 77 | 990 | 0.62 |
| Y | 14 | 200 | 1,975 | 9.9 | 600 | 4,700 | 0.71 |
| Y | 8 | 2,000 | 7,328 | 3.7 | 4,300 | 18,000 | 0.61 |
| Y | 8 | 2,000 | 6,250 | 3.1 | 2,500 | 16,000 | 0.70 |
| Y | 8 | 2,000 | 3,338 | 1.7 | 2,000 | 4,500 | 0.29 |
| Y | 14 | 2,000 | 2,121 | 1.1 | 570 | 4,518 | 0.48 |

